# Intense expression of Neurokinin-1 Receptor is associated with Urothelial carcinoma

**DOI:** 10.1101/2020.06.27.175026

**Authors:** Riffat Mehboob, Amber Hassan, Shahida Perveen, Syed Amir Gilani, Humera Waseem, Fridoon Jawad Ahmad, Javed Akram

## Abstract

**I+ntroduction:** Substance P (SP)/ Neurokinin-1 Receptor (NK-1R), induces inflammatory reactions in peripheral tissues but its regulatory effects in target tissues are dependent on receptor signaling. SP has a high affinity for the NK-1 receptor, to which it binds preferentially. SP/NK-1R complex plays a key role in the interaction in the onset of pain and inflammation.

**Objective:** The aim of this study was to investigate the expression of NK-1R in Urotheilial carcinoma and its association with cancer progression.

**Method:** The study included ten biopsy samples of Urinary bladder, obtained retrospectively from a tertiary care hospital of Lahore. An Immunohistochemical study was performed using monoclonal antibodies against NK-1R. The presence or absence of staining and the intensity of the immunoreactivity were noted, as well as the number and type of cells. Evaluation of the Immunohistochemical expression was determined by the semi-quantitative method and scored.

**Result:** NK-1 receptor was intensely expressed in patients with higher grade Urotheilial carcinoma. The cases clinically diagnosed as High Grade Urotheilial Carcinoma showed intense expression of NK-1R. However, the cases clinically diagnosed as low grade Urotheilial carcinoma showed very weak staining with NK-1R. However, the normal margins of the same tissue showed negative expression.

**Conclusions:** Elevated expression of NK-1R was associated with advanced stage of urothelial carcinoma. It is the first study to our knowledge to report this association. It may serve as a good diagnostic as well as prognostic marker and therapeutic strategy.

## 1.1. INTRODUCTION

Neuropeptides have been implicated in inflammation in peripheral tissues as well as modulation of pain processes. These peptides include Substance P (SP), calcitonin gene-related peptide (CGRP), neurokinin A, neuropeptide Y, and vasoactive intestinal polypeptide (VIP) among others.^1^

Prior studies have demonstrated SP to be involved in both inflammation^2^ and pain^3^. Levels of SP released by the sensory fibers is further enhanced as a consequence of inflammation and this viscious circle continues.^3^ All the biological actions of SP are exerted by binding to G-protein-coupled receptor, NK-1 which is located on most of the inflammatory cells e.g. mast cells, macrophages and other immune cells.^4^

The interaction of SP with mast cells describes the reason for the enhanced vascular permeability and blood pressure due to induction of release of histamine. Moreover, macrophages lymphocytes and granulocytes, contain sites of NK-1R, and these cells are stimulated by SP and induce the production of proinflammatory mediators and cytokines.^5^

Few studies have reported the involvement of SP and NK-1R with the progression of different carcinomas^2, 6-10^ including our own previous study^11^. However, there is no evidence of the behaviour of NK-1R expression during normal and inflamed stages in human Urotheilial carcinoma. It will be the first study to evaluate the expression of NK-1R in urothelial carcinoma. This study was thus aimed to investigate the expression of NK-1R in Urotheilial carcinoma and its association with cancer progression.

## 1.2. MATERIAL AND METHODS

### 1.2.1. Tissue Collection

The protocol and procedure of the study were performed after taking the Ethical approval from the Ethical Committee of the University of Lahore, Lahore, Pakistan. Ten biopsy samples of urinary bladder were obtained from a tertiary care hospital of Lahore. Histopathological and immunohistochemical evaluation was performed at Research Unit, Faculty of Allied Health Sciences, The University of Lahore.

### 1.2.2. Immunohistochemistry for SP and NK-1R

The section of 3 to 5 µm was cut from the paraffin embedded tissue block of urinary bladder for the immunohistopathological detection of NK-1 receptors. After the overnight incubation at 4°C NK-1R (Abbott, 1:1000) staining was performed manually. Sections were incubated in peroxidase-blocking solution and heated at 100 °C for 60 min. After incubation for 32 min at 37 °C, the tissue sections were incubated with a universal secondary antibody (Roche Diagnostics KK) for 20 min at 37 °C and then visualized by the DAB Map detection kit (Roche Diagnostics KK). As the negative control the primary antibody was omitted and was being replaced by the non-immune serum. Nuclei were counterstained using a hematoxylin counterstain reagent (Roche Diagnostics KK). We performed an antigen retrieval step in 10 mM citrate buffer (pH 6.0) before the immune-histochemical staining. We studied representative samples of UB. The evaluated slides were evaluated by two independent pathologists. In each slide, high-power microscopic fields were evaluated using a 40X magnification.

The presence or absence of staining and the intensity of the immunoreactivity were noted, as well as the number and type of cells showing a brown staining and whether or not the staining was localized in the nucleus, cytoplasm cells and/or in the plasma membrane. The results were recorded as positive when they showed cellular and/or plasma membrane staining ranging from moderate to strong in more than 10% of the cells. By consensus among the pathologists, the intensity of the immune-reactive cells was scored as follows: when less than 10% of the total cells were stained, the number of immune-reactive cells was considered low, it was considered moderate when 10-40% were stained and high when more than 40% were stained^11^. The specimens were examined and photo-graphed at 10, 20 and 40X magnification utilizing a digital microscope camera (Olympus AX80 DP21; Olympus, Tokyo, Japan) interfaced with a computer. All protein levels were evaluated using the nuclear labeling index (%), recorded as the percentage of positively stained nuclei in 100 cells in the hot spot.

### 1.2.3. Statistical Analysis

Data are presented as mean percentages of positive cells ± standard deviation. The correlation between the nuclear labeling indexes was assessed by Spearman’s rank correlation. *p* < 0.05 was considered to indicate statistical significance. All data analyses were carried out using the Statistical Package for Social Sciences, version 20 (SPSS Inc., Chicago, IL, USA).

## 1.3. Results

The mean age of patients was 64.8 ± 5.73 years. There were 7 (70.0%) males and 3 (30.0%) females, negative NK-1R staining was observed in 1(10.0%), mild in 3 ((30.0%) and severe in 6 (60.0%) cases. 4 (40.0%) patients had low grade carcinoma and 6 (60.0%) patients had high grade carcinoma (Table 1, Figure 1). There was strong positive correlation between the grades of carcinoma and NK-1R expression. NK-1R expression and intensity increased with advancement in grading (0.976**, p-value 0.000) (Table 2, Figure 2).

**Table 1:** Neurokinin-1 Receptor expression and Grading of carcinoma.

| Variables | Frequency | Percentage (%) |
| --- | --- | --- |
| **Intensity of NK-1R Staining** |  |  |
| **Negative** | 1 | 10 |
| **Mild** | 3 | 30 |
| **Severe** | 6 | 60 |
| **Grade of Cancer** |  |  |
| **Low Grade Cancer** | 4 | 40 |
| **High Grade Cancer** | 6 | 60 |

**Table 2:**
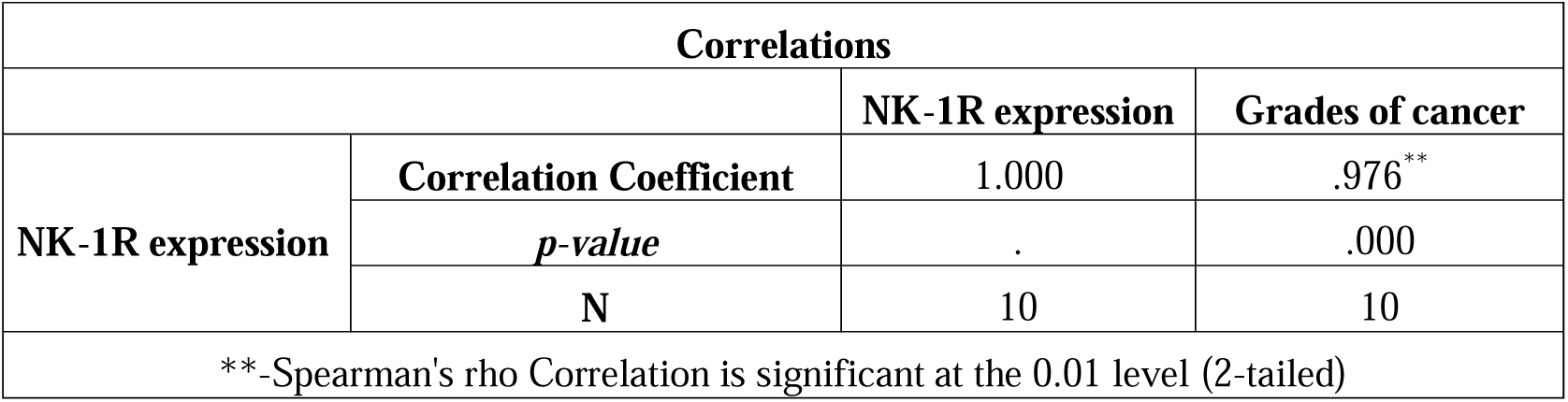
Spearman correlation.

**Figure 1-A).**
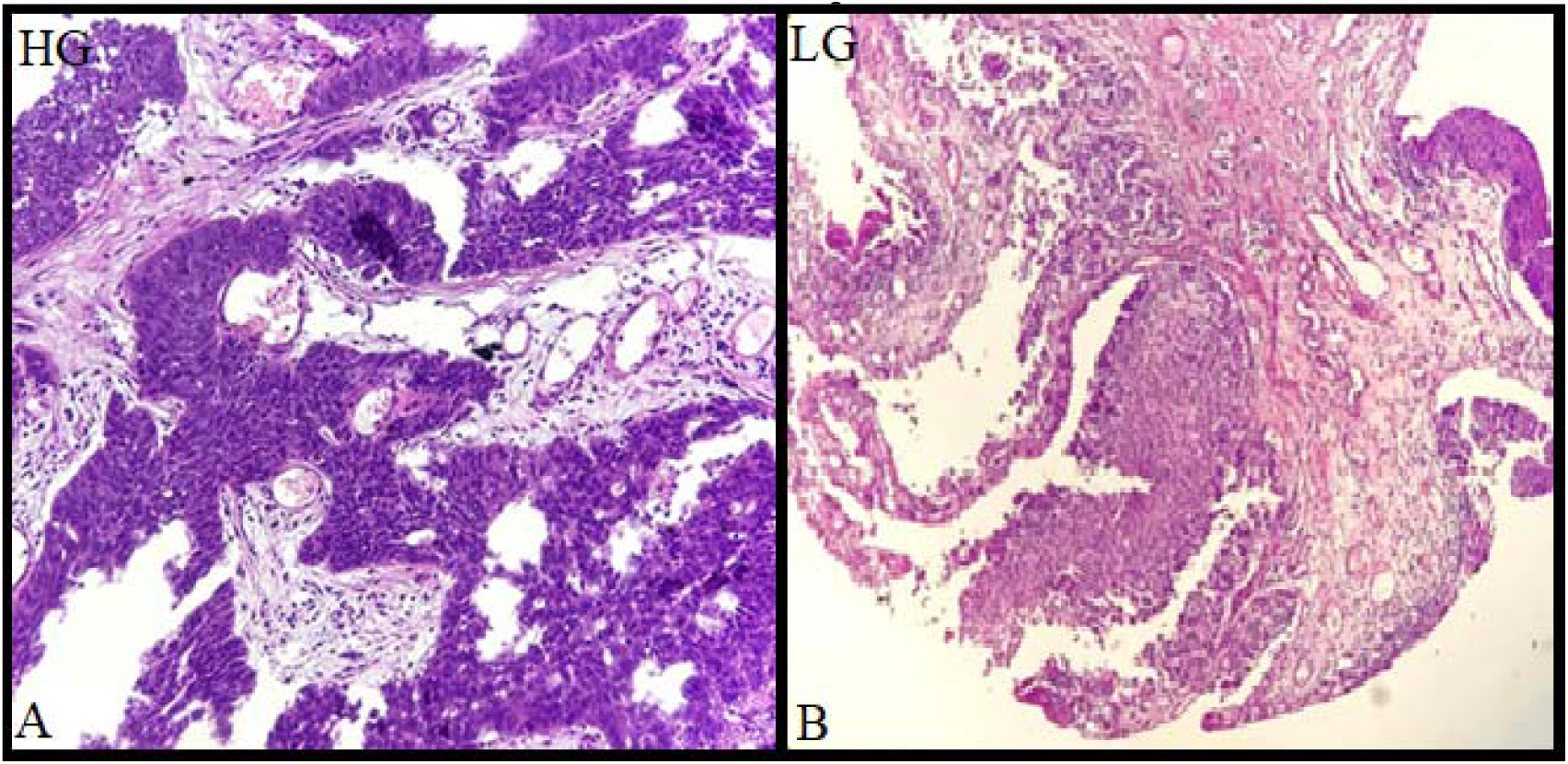
Photomicrograph of high grade UC (Transitional cell carcinoma) with groups and sheets of neoplastic cells with coarse chromatin open up nuclei, 20X; B) Photomicrograph of low grade UC (Transitional cell carcinoma) with more than 10 layers thick neoplastic cells infiltrating the lamina propria, 20X-HG-High grade; LG-Low grade

**Figure 2-A).**
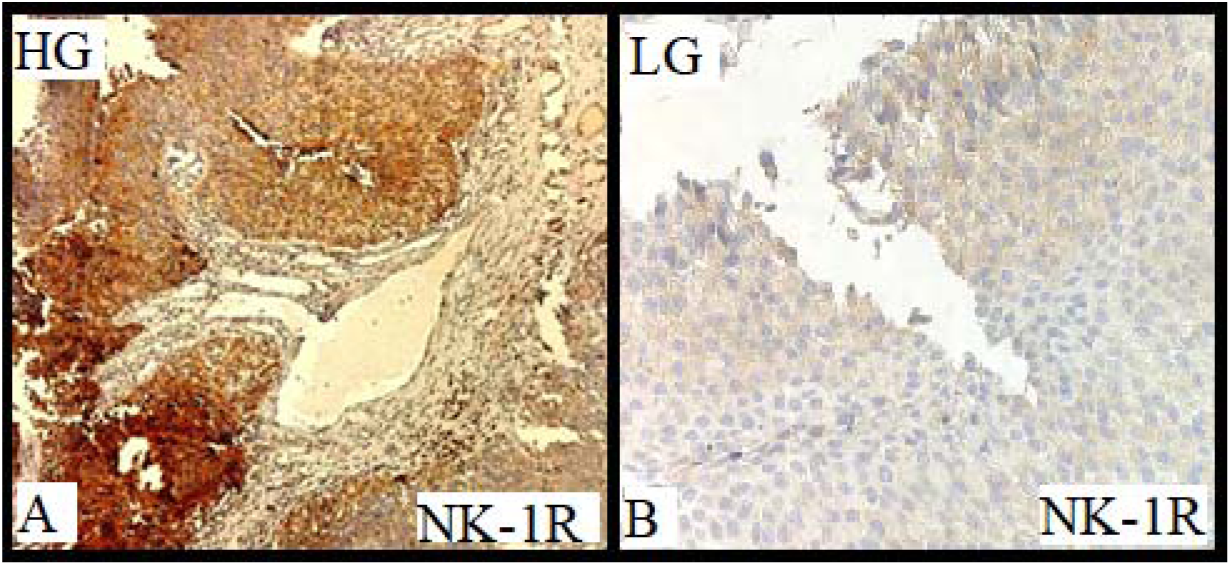
High grade Urothelial carcinoma with strong positive stain For NK-1R; B) Low grade Urothelial carcinoma with weak NK-1R staining

## 1.4. Discussion

SP/ NK-1R induces inflammatory reactions in peripheral tissues but its regulatory effects in target tissues are dependent on receptor signalling.^13^ The objective of this study was, therefore, to determine the NK-1R expression in urinary bladder tissue having a clinical diagnosis of Urothelial carcinoma. The mechanism by which inflammation arises when an inflammatory reaction is going on within the cancer progression is not completely understood. This study proved that NK-1R expression was found in these samples and there was an intense expression in high grade cases. One possible mechanism may be the increased secretion of SP from trigeminal ganglion in response to any inflammation or nociceptive stimulus. This SP is not only neurotransmitter but neuromodulator, neurohormone and immunomodulatory. It will bind to its receptor NK-1R in the target cells. Under normal physiological conditions, it is present in brain and gut. But in pathological conditions its increased expression may cause cancer^6, 8, 12^, sudden death ^13-16^ as well as pain^3^.

This study is in agreement with the previous studies reporting the association of SP/ NK-1R mechanisms in cancer progression ^7, 11^ and suggest its therapeutic role along with diagnostic and prognostic one. This is the first study to report the expression of NK-1R in Urotheilial carcinoma and its association with its progression. These findings have clinical as well as diagnostic and therapeutic significance. The identification of involvement of NK-1R as a pathological mechanism in human urothelial carcinoma and its association with disease progression is important for widening of understanding regarding the mode of action. It may render a good therapeutic option as well as management of such patients.

## 1.5. Conclusions

NK-1R is expressed in urothelial carcinoma with a significantly increased expression during clinical advanced stage. This study may be considered as a pioneer study to show the expression of Neurokinin-1 Receptor and its higher intensity in advanced stage of carcinoma. However, it should be explored further on larger sample size. Its antagonists may serve as a therapeutic regimen for the treatment of bladder carcinoma. It may also serve as a diagnostic marker to differentiate the stages as well as to diagnose the carcinoma.

